# Dissociable Spatial and Feature Tuning of Gamma and Alpha Rhythms in Human Visual Cortex

**DOI:** 10.1101/2025.08.09.669461

**Authors:** Sanaz Ghaffari, Arian Yavari, Sara Bonyadian, Arsalan Ghofrani, Russell Butler

**Affiliations:** Bishop’s University

## Abstract

Visual stimulation in humans reliably induces narrowband gamma (40–80 Hz) increases and alpha (8–12 Hz) suppression in EEG, but the relationship between these rhythms is not fixed. Using high-density EEG, we mapped gamma and alpha responses across a broad set of retinotopic, orientation, and motion conditions. Gamma was strongly retinotopically tuned, with linear summation of subfield responses accurately predicting full-field responses. In contrast, alpha showed little spatial tuning, substantial responses even in the absence of visual input (anticipatory suppression), and subadditive summation consistent with divisive normalization. Across most retinotopic configurations, higher gamma coincided with stronger alpha suppression, yet systematic dissociations emerged, with full-field and foveal-centered gratings evoking stronger than expected gamma relative to alpha suppression. Orientation tuning was robust for gamma but weak for alpha, with oblique gratings producing high gamma yet weak alpha suppression, reversing the usual inverse relationship. These patterns indicate that EEG gamma power primarily reflects large-scale synchronous activity that matches the ‘global LFP’ observed in macaque V1, whose spatial and feature tuning properties are independent of both single-neuron selectivity and feedback-driven alpha dynamics. The results establish a mechanistic dissociation between gamma and alpha rhythms, highlighting distinct circuit origins and tuning principles for these canonical visual responses.

## Introduction

Gamma (∼40–80 Hz) and alpha (∼8–12 Hz) oscillations are prominent visual cortex rhythms with opposing functional associations. Alpha, first described by Berger in 1929 [1] and later characterized by Adrian and Matthews [2], is high during eyes-closed rest and suppresses upon visual stimulation (“blocking”), reflecting an idling or inhibitory state. Gamma, discovered later in animal visual cortex [3], emerges during active visual processing, reflecting synchronous excitatory–inhibitory interactions. Human EEG, MEG, and ECoG studies confirm that stimulus-induced gamma is linked to feature encoding, perceptual binding, and attention [4]. Thus, gamma generally marks active processing, whereas alpha marks relative inactivity or inhibition [11].

Their tuning and spatial profiles differ sharply. Gamma is localized, retinotopically specific, and sharply tuned to features such as orientation, spatial frequency, and motion direction, sometimes mirroring local spiking preferences [5,6,9]. For instance, optimal-orientation gratings evoke strong gamma, whereas orthogonal orientations do not [5]. Alpha shows weak feature tuning: most salient visual stimuli suppress alpha regardless of specific features, with only modest orientation dependence [11]. While alpha suppression can be region-specific (e.g., over stimulated field representations), its spatial focus is broad compared to gamma [8]. In sum, gamma reflects local, feature-selective excitation, whereas alpha reflects diffuse inhibitory control, generally inversely related to visual drive. Gamma and alpha oscillations appear to mediate complementary roles in cortical communication. Converging evidence links gamma to feedforward sensory signaling and alpha to feedback modulation and inhibition [7,8]. For example, van Kerkoerle et al. [7] showed that visual stimulation drives gamma feedforward from V1 to higher areas, while alpha travels as feedback from higher to lower areas. Microstimulation of V4 induced alpha in V1, whereas stimulating V1 evoked gamma in V4.

Although the feedforward–feedback framework provides a useful starting point, important gaps remain in understanding how gamma and alpha relate in the human visual cortex. It is still unclear whether they emerge from a single push–pull circuit or from largely independent neural generators, perhaps in different cortical layers or cell populations. Nor is it known whether their apparent antagonism is obligatory—whether increases in one rhythm necessarily drive decreases in the other—or if they can vary independently under certain stimulus or cognitive conditions.

One underexplored dimension concerns how spatial integration shapes these rhythms. In primate V1, large visual stimuli can synchronize local gamma into a coherent “global gamma” spanning millimeters of cortex [6,9,10], a process thought to explain the strong gamma “bump” observed in human EEG/MEG for large-field stimuli. Whether this synchronization is accompanied by proportionally broader alpha suppression, or whether alpha responds in a more context-dependent manner, is not well established. Furthermore, most prior work has studied gamma and alpha tuning in isolation, leaving open the question of how they behave when measured in parallel under identical stimulus manipulations—particularly across both spatial and feature dimensions.

To address these questions, we directly compared the spatial and feature tuning of gamma and alpha oscillations in human visual cortex using EEG. By systematically varying retinotopic location, stimulus size, orientation, and motion direction—while also including no-stimulus baseline conditions—we tested whether gamma and alpha follow a shared tuning logic or instead reflect distinct encoding mechanisms. This approach allowed us to probe whether gamma’s spatial specificity and feature selectivity are mirrored by alpha, to quantify how anticipatory alpha suppression influences spatial summation, and to reveal circumstances in which gamma–alpha relationships deviate from their typical inverse correlation. The results, summarized below, provide new evidence for a tight but non-uniform coupling between feedforward-driven gamma synchronization and feedback-mediated alpha modulation.

## Materials and Methods

**Subjects:**thirty healthy young adults (17 male, 13 female; age range: 18–38 years) were recruited from the Bishop’s University and Université de Sherbrooke communities. All participants provided written informed consent in accordance with ethics approval from the *Centre intégré universitaire de santé et de services sociaux de l’Estrie – Centre hospitalier universitaire de Sherbrooke (CIUSSS de l’Estrie – CHUS)*. All methods were performed in accordance with the relevant guidelines and regulations. Participants were screened to exclude any neurological or psychiatric disorders, sleep disorders, or use of medications affecting the central nervous system.

### Stimulus Presentation

Stimuli were generated in MATLAB (The MathWorks, Inc., Natick, MA) using Psychophysics Toolbox extensions and presented on a 15-inch CRT monitor in a dimly lit, sound-attenuated room. The display background was uniform gray, with luminance matched to the mean luminance of each stimulus. The grating stimuli (both bars and annulus) had a spatial frequency of 3 cycles/° and a temporal frequency of 6 cycles/s, the full field stimulus had a aperture width of 10° diameter centered on fixation.

The stimulus set consisted of 36 distinct trigger types (see Figure 1):

**Figure 1.**
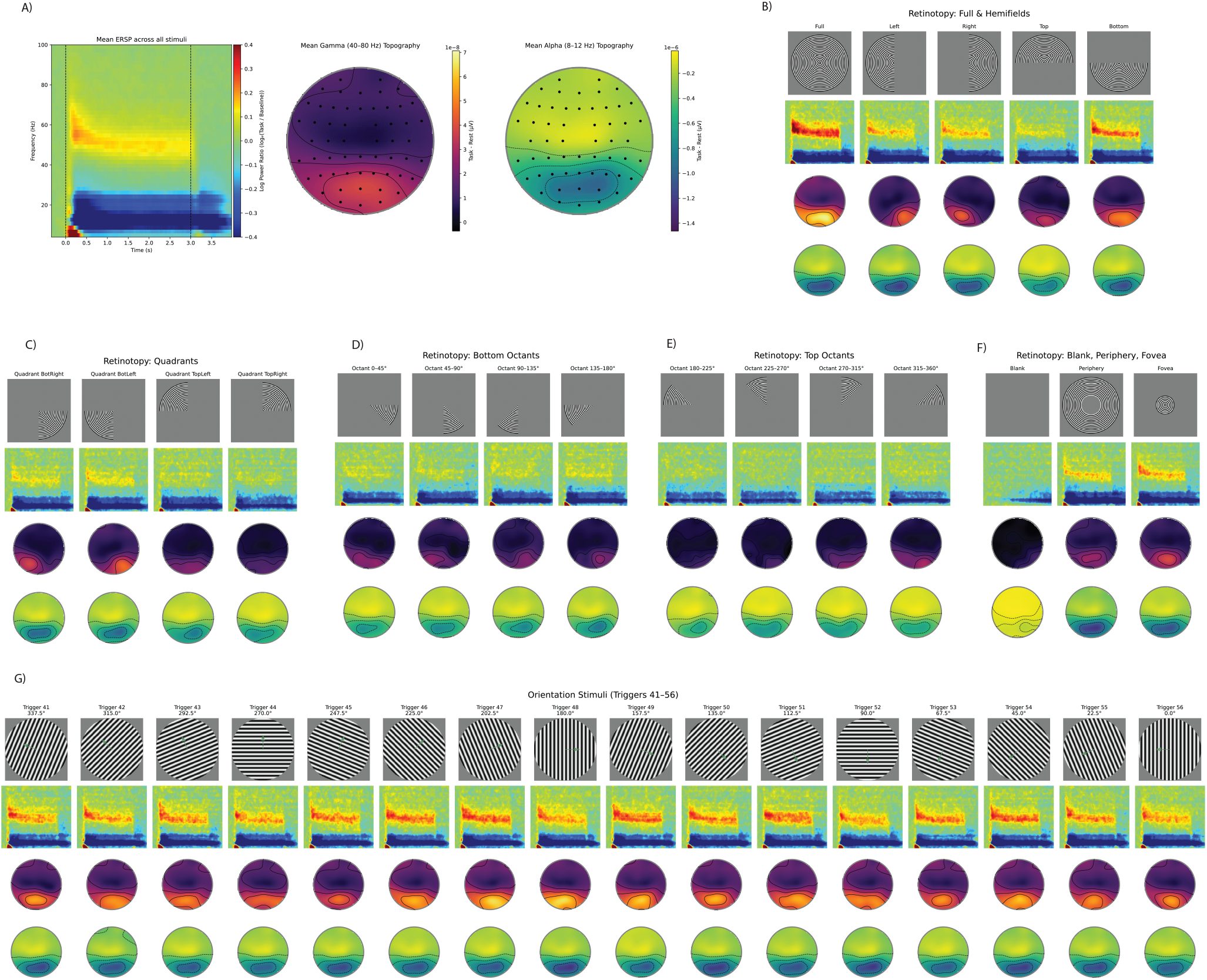
Raw data and qualitative results. (A) Time–frequency plots and topographic maps for gamma (40–100 Hz) and alpha (8–12 Hz) bands. (B) Topographic maps for full-eld and hemi eld stimuli. (C) Topographic maps for quadrant stimuli. (D) Topographic maps for octant stimuli (0°–180°). (E) Topographic maps for octant stimuli (180°–360°). (F) Topographic maps for blank, foveal, and peripheral stimuli. (G) Time–frequency plots and topographic maps for 16 oriented grating stimuli.

**Figure 2.**
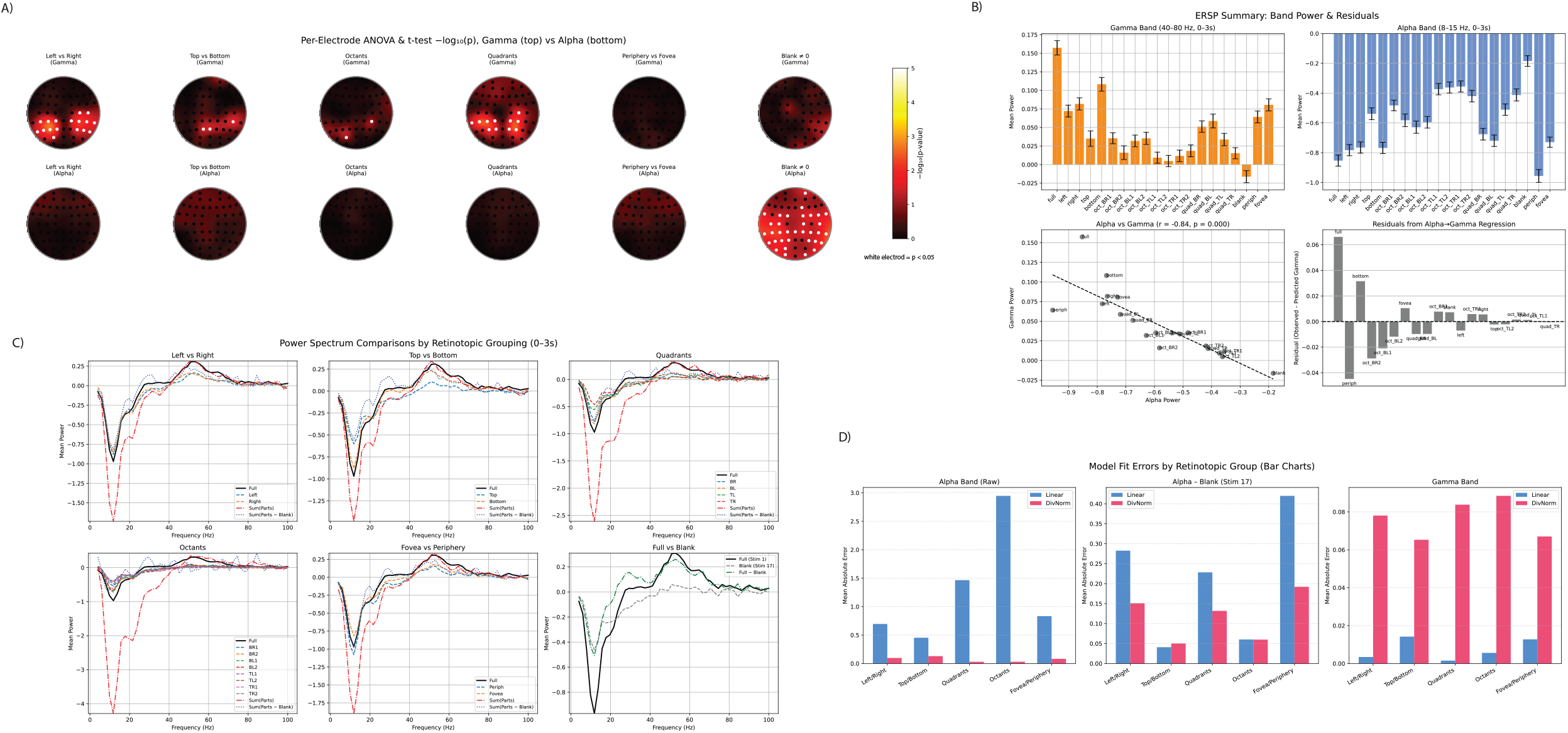
Retinotopic Tuning of Induced Gamma and Alpha Oscillations. (A) Scalp-wise –log10(p) maps from ANOVAs and t-tests for gamma (top row) and alpha (bottom row) power across retinotopic groupings, including left/right, top/bottom, quadrants, octants, and blank. (B) Band-power summaries (0–3 s) for gamma (top left) and alpha (top right) across conditions; scatterplot of alpha vs. gamma power (bottom left); regression residuals (bottom right). (C) Full-eld spectra vs. summed sub eld responses for gamma and alpha, with and without blank correction, across retinotopic groupings. (D) Model prediction errors for linear vs. divisive normalization ts, with and without blank correction, for alpha (left, center) and gamma (right) bands.

**Figure 3.**
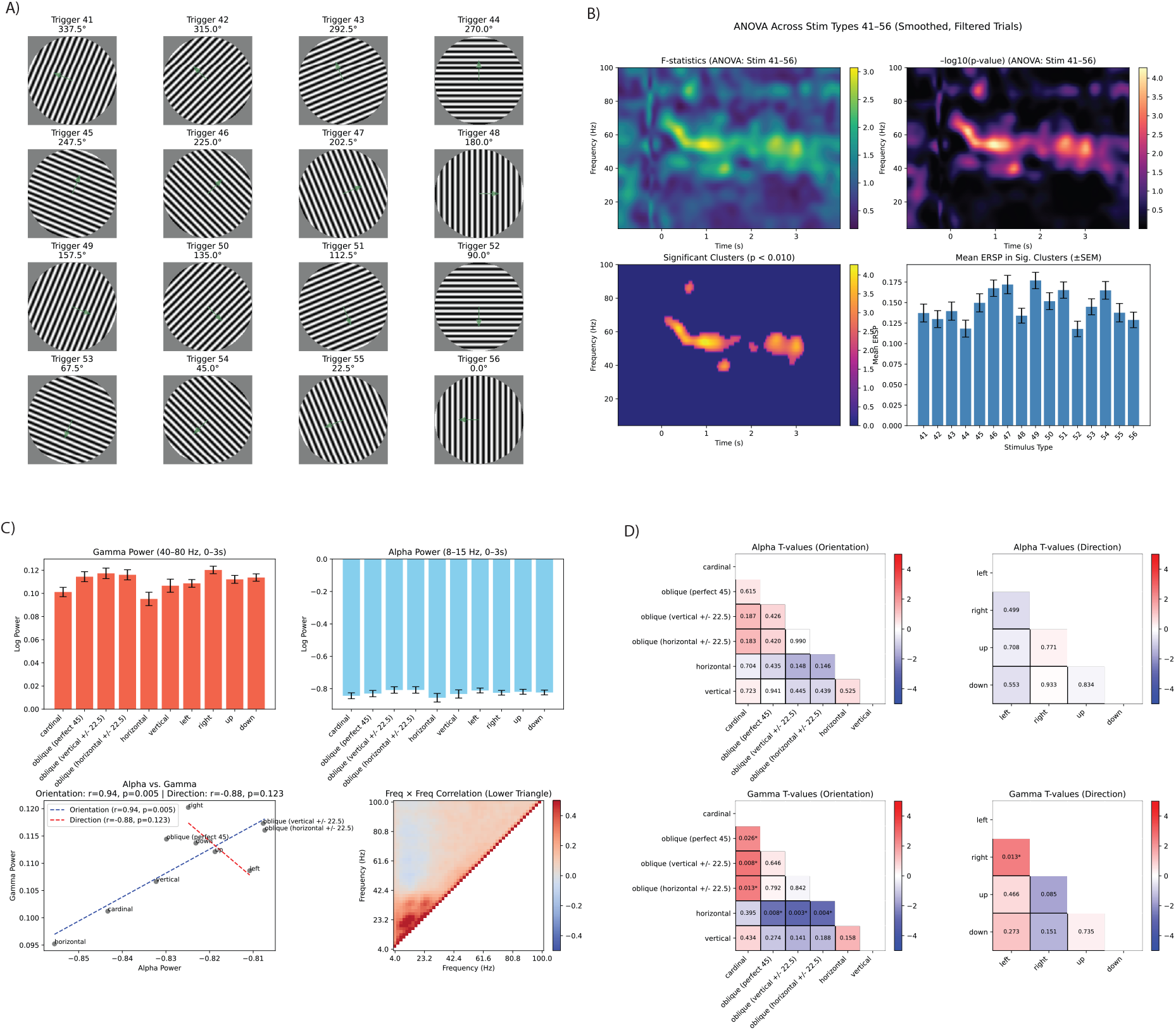
Orientation tuning. (A) Sixteen oriented grating stimuli (Triggers 41–56) spaced at 22.5° intervals from 0° to 360°. (B) One-way repeated-measures ANOVA results: F-statistic map (top left), –log10(p) map (top right), cluster-based permutation test (bottom left), and mean gamma-band ERSPs for all orientations (bottom right). (C) Band-power summaries (0–3 s) for gamma (top left) and alpha (top right) across orientations; scatterplots for gamma vs. alpha (bottom left) and drift direction groupings; frequency–frequency correlation matrix (bottom right). (D) Pairwise t-tests for orientation (left column) and drift direction (right column) e ects on alpha (top row) and gamma (bottom row) power.

- **Retinotopic mapping stimuli**: full-field; left/right hemifields; upper/lower hemifields; four quadrants; eight octants; central fovea; peripheral ring; and a blank “anticipatory” condition with no visual stimulus. Retinotopic masks were applied to a 100% contrast inward-drifting circular annulus.
- **Orientation stimuli**: 16 high-contrast drifting gratings, starting at vertical (0°) drifting leftward, and rotated in 22.5° increments through 337.5°. Motion was always orthogonal to grating orientation.

All stimuli were presented at 100% Michelson contrast against the gray background. Retinotopic stimuli were spatially masked using binary apertures smoothed with a Gaussian edge, while orientation stimuli were unmasked except for the circular aperture.

Each trial began with a **fixation period** in which a black crosshair was presented. Immediately before stimulus onset, the crosshair turned red for a randomized foreperiod of 0.25 ± 0.15 s, serving as a temporal cue that the stimulus would appear. This red “primer” was also used before blank trials in the anticipatory condition. Stimuli were then displayed for 3 s, followed by a 1 s post-stimulus fixation interval before the next trial began.

Stimuli were presented in ‘batches’ of 8 minutes, with participants maintaining central fixation throughout each batch and given1-2 minute rest periods between. Stimuli were presented in pseudorandom order, with 10–20 repetitions per stimulus type (360–720 total trials per participant). The total session lasted approximately 1 hour.

### EEG Preprocessing – Independent Component Analysis (ICA) Extraction

Continuous EEG data were acquired in BrainVision format (.vhdr) and preprocessed using MNE-Python (v1.8.0) with the Picard ICA algorithm (Picard v0.8). All preprocessing was performed in Python (v3.9.21).

Each.vhdr file was imported using mne.io.read_raw_brainvision() with data preloaded into memory. To identify bad channels, the raw signal was band-pass filtered between 10–100 Hz (FIR filter, firwin design) and the sum of squared differences (SSD) was computed for each channel. Z-scores of SSD values were calculated, and channels with *z* > 1 were marked as bad and subsequently interpolated using spherical spline interpolation, resulting the removal of approximately 1-3 channels per subject.

Line noise was removed using a 60 Hz notch filter, and a copy of the data was then band-pass filtered between 1–100 Hz (firwin design) for ICA. This filtered dataset was resampled to 200 Hz to reduce computational load. Events were extracted from annotations, and epochs were defined from ࢤ1.0 s to +4.0 s relative to event onsets and concatenated into a continuous Raw object for ICA training.

ICA was performed using the Picard algorithm with n_components=60, extended=True, and a maximum of 500 iterations. The ICA model was fit to the concatenated epochs, and the resulting decomposition (unmixing and mixing matrices, component activations) was saved in.fif format for later use in artifact identification and removal.

### Artifact identification and removal

For each component, the mean ERSP across trials was visualized alongside its corresponding scalp topography derived from the ICA weight matrix. Components were manually inspected and classified based on both their spectral and spatial profiles. Components showing narrowband gamma activity (30–100 Hz) with posterior scalp topographies, as well as those with narrowband alpha activity (8–12 Hz), were retained for further analysis. All selected “good” components were saved in subject-specific files for use in subsequent stages of signal reconstruction and analysis, typically 3-5 components were retained per subject.

The saved ICA solution was then used to extract source activations for only the retained components. Event-locked epochs (−1.6 s to +4.6 s, including a 0.6 s buffer) were segmented from these sources for all stimulus triggers of interest. Time–frequency representations (4–100 Hz) were computed using Morlet wavelet convolution, cropped to the analysis window (−1.0 s to +4.0 s) to remove padding, and log-transformed. Single-trial baseline correction was performed for each component and frequency bin using the mean power from −0.5 s to 0 s pre-stimulus, producing baseline-normalized ERSPs for every trial, component, and frequency–time bin. These single-trial ERSPs were then stored for each subject along with their corresponding stimulus IDs, enabling both subject-level and stimulus-level averaging in later analyses.

### Orientation Tuning Analysis

Orientation tuning analyses were performed on baseline-corrected, single-trial ERSPs from all retained ICA components across subjects. To isolate trials corrupted by broadband muscular or ocular artifacts, we first extracted log power in the 80–100 Hz band (broadband gamma) for each trial. The maximum power within the analysis window (0–3 s) was computed per trial, and values were z-scored across all subjects. Trials with |z| > 4 were excluded to remove extreme outliers (the same technique was applied in retinotopy analysis).

To assess orientation selectivity, a two-dimensional repeated-measures approach was used. First, all single-trial ERSPs were Gaussian-smoothed (σ = 2 freq bins, 4 time bins) to increase spatial coherence. A pointwise one-way ANOVA (factor: stimulus identity) was then conducted across all grating orientations for each frequency–time bin, yielding F- and p-value maps. p-values were transformed to –log□□scale, and significant clusters (p < 0.01, cluster-based spatial connectivity) were identified using an 8-connected neighborhood criterion. The size and location of significant clusters were visualized, and mean ERSP values within these clusters were extracted per stimulus type. Cluster-averaged ERSPs were compared across orientations using bar plots (mean ± SEM).

Stimuli were further binned into predefined orientation and motion-direction categories (e.g., “cardinal,” “oblique ± 22.5°,” “vertical,” “horizontal,” “left,” “right,” “up,” “down”). Within each bin, mean gamma power (40–80 Hz, 0–3 s) and alpha power (8–15 Hz, 0–3 s) were computed, along with standard errors. Pearson correlations between bin-averaged alpha and gamma power were calculated separately for orientation bins and for direction bins, and linear fits were overlaid on scatter plots with correlation coefficients and p-values reported.

To examine pairwise differences between bins, two-sample Welch’s t-tests were performed on the trial-level alpha and gamma values for each bin combination. This produced t-value and p-value matrices for both orientation and direction conditions, which were visualized as lower-triangle heatmaps with p-values displayed in each cell and significance denoted by an asterisk (p < 0.05). Finally, a frequency × frequency correlation matrix (Pearson’s r) was computed from the mean post-stimulus (0–3 s) power spectra across all trials.

### Computation of Per-Electrode Power

For each subject, cleaned EEG data were reconstructed in sensor space by loading the subject’s saved ICA decomposition, marking all components not in the good-component list for exclusion, and applying the ICA to the continuous EEG, thereby removing artifactual sources while retaining physiologically meaningful activity. The cleaned scalp-space EEG was segmented into epochs time-locked to each stimulus (−0.5 to 3.5 s), and narrowband alpha (8–13 Hz) and gamma (40–80 Hz) signals were isolated by applying zero-phase FIR band-pass filters to separate copies of the data. For each trial, absolute analytic amplitude was computed at every electrode, and the mean amplitude in the stimulus window (0–3.0 s) was compared to the mean amplitude in the pre-stimulus baseline (−0.5–0 s). The final power measure at each electrode was the difference between these two means, expressed as |task| – |baseline|. This computation was performed separately for alpha and gamma, averaged across trials within each stimulus type, and stored both as trial-level arrays and as per-stimulus, per-channel matrices.

### Topographic Visualization of Alpha and Gamma Power

To display spatial distributions of stimulus-evoked power, electrode coordinates were extracted from a representative subject’s EEG montage and projected to 2-D using spherical coordinates. Channels with poor coverage or located at the extreme periphery (e.g., mastoid, inferior temporal) were excluded to improve interpolation stability. For each stimulus type, the mean alpha or gamma difference values across subjects were interpolated over a 2-D grid using radial basis functions (multiquadric kernel, smooth = 0.1). The interpolation grid was circularly masked to match the scalp outline, with a slightly expanded radius to ensure outermost electrodes were represented.

### Retinotopy analysis

Retinotopy figures were generated from the same baseline-corrected, single-trial ERSP datasets described earlier, using all retained ICA components. Band-specific power measures were computed by averaging over both time (0–3 s) and the relevant frequency range—8–15 Hz for alpha, 40–80 Hz for gamma. Mean and SEM values were calculated for each stimulus, and results were summarized in bar plots for each band. Alpha and gamma means were then compared directly in a scatter plot with a least-squares regression fit, reporting correlation coefficients and p-values, and residuals from the regression were plotted to highlight stimuli deviating most from the alpha–gamma relationship.

### Retinotopic Group Comparisons and Model Fits

To examine spatial summation properties, stimuli were grouped into standard retinotopy configurations: left vs right hemifields, top vs bottom hemifields, quadrants, octants, and fovea vs periphery. For each grouping, the full-field stimulus spectrum was compared to (a) the spectra for each part stimulus, (b) the sum of the part spectra, and (c) the sum of the part spectra with the blank-field (no stimulus) spectrum subtracted. These were overlaid in line plots to visualize how closely part-sum responses matched the full-field response.

Finally, quantitative model fitting compared two summation models—a simple linear sum and a divisive normalization (DivNorm) model with fixed σ = 0.5—applied separately to alpha and gamma responses. For each grouping, mean absolute errors between model predictions and the observed full-field response were computed, with an additional variant for alpha in which the mean blank stimulus power was subtracted prior to fitting. These errors were summarized in grouped bar charts for each band and model type, allowing direct visual comparison of linear versus normalized summation performance across retinotopic configurations.

### Per-Electrode Statistical Mapping of Retinotopic Effects

Per-electrode one-way ANOVA was performed across the grouped conditions; for the special “Blank ≠ 0” test, a per-electrode one-sample t-test against zero was used. The resulting p-values were converted to −log□□(p) for visualization.

## Results

The spectral and topographical analyses in Panel A revealed robust stimulus-induced gamma-band (40–100 Hz) increases and alpha-band (8–12 Hz) decreases following stimulus onset across all trials. These responses were temporally aligned with visual presentation (time 0) and showed distinct spatial distributions. Gamma-band power was maximal over occipital electrodes, indicating localized visual cortex activation, while alpha suppression extended broadly across posterior scalp regions, consistent with desynchronization during visual engagement. The mean ERSP plot (left) confirms a prominent increase in gamma power and concurrent alpha suppression, with alpha suppression also evident in the post-stimulus window (3 – 3.5 s).

Panels B–F qualitatively demonstrate the spatial specificity of induced oscillations across retinotopic stimuli. Full-field and hemifield stimulation (Panel B) elicited the strongest gamma responses for full and lower field stimuli, consistent with the anatomical overrepresentation of the lower visual field in primary visual cortex, and left/right gratings evoking contralateral gamma in the scalp maps. In Panel C, quadrant stimuli produced distinct topographic gamma patterns: lower-left and lower-right quadrants showed strong focal activation in contralateral occipital areas, whereas upper quadrant responses were weaker and more diffuse. Panels D and E further refine these findings with octant-level resolution, showing that bottom-octant stimuli consistently produced the largest gamma increases, especially between 90°–135° and 135°–180°, whereas top-octant responses were notably weaker. Panel F revealed that the “Blank” condition evoked negligible gamma power but a strong delayed and post-stimulus alpha response, while peripheral stimulation activated lateral occipital electrodes and central (foveal) stimulation elicited more focal midline occipital gamma. Qualitatively, alpha responses were less focal and retinotopically tuned than gamma responses.

Finally, Panel G displayed gamma responses across 16 oriented grating stimuli. All orientations produced a similar temporal profile of induced gamma, but notable topographic variability was observed across orientations. Certain angles (e.g., 45°, 90°, 135°) produced more focal or intense gamma activity than others, hinting at potential orientation tuning or bias in the cortical response. Importantly, alpha suppression remained relatively consistent across conditions, affirming its role as a general marker of visual engagement rather than stimulus-specific processing. Together, these findings demonstrate spatially specific gamma-band activation tied to retinotopic and orientation features, underpinned by a global alpha suppression signature of visual processing.

Panel A displays the scalp-wise –log10(p) maps from ANOVAs and t-tests comparing per-electrode gamma (top row) and alpha (bottom row) power across multiple retinotopic groupings. Gamma responses showed robust spatial selectivity, with significant electrode clusters (p < 0.01, bright red/yellow) in occipital regions differentiating left vs right, top vs bottom, and full retinotopic subdivisions such as quadrants and octants. In contrast, alpha showed no or very limited spatial selectivity, with none of the electrodes showing a significant difference in alpha power due to retinotopic configuration. The last column (Blank ≠ 0) shows that the gamma response to a plain gray background was not significantly elevated in any electrode, while alpha shows widespread and diffuse suppression even in response to the null stimulus, matching the non-spatial selectivity shown by alpha in the first 5 columns.

Panel B summarizes induced band power (0–3 s) across all retinotopic conditions based on the ERSP averaged across all components (pooling responses across all posterior electrodes). Gamma power (top left) was highest for stimuli covering larger or more complex visual areas (e.g., “full” and “bottom”), and weaker for peripheral or upper-field stimuli. Alpha suppression (top right) followed a similar pattern, with the more alpha suppression in response to full-field or bottom hemifield stimuli. Notably, the upper-field octants such as TL1 and TL2 (top-left 1 and 2) still showed robust alpha suppression despite a complete lack of gamma response. This relationship was quantified in a scatterplot of alpha vs. gamma power across conditions (bottom left) revealed a strong inverse correlation (r = –0.84, p < 0.0001), reinforcing the antagonistic relationship between these rhythms. However, residuals from the regression (bottom right) highlighted notable deviations, such as enhanced gamma (relative to alpha) for “bottom” and “full” field stimuli, and suppressed gamma for the ‘peripheral’ grating beyond what alpha suppression alone would predict.

Panel C compares full-field power spectra to the summed responses of spatially distinct retinotopic subregions, revealing clear differences in spectral linearity between frequency bands. Across all comparisons—left vs right, top vs bottom, quadrants, octants, and fovea vs periphery—the summed gamma-band responses (40–80 Hz, red dashed line) closely approximate the full-field response (black solid line), indicating near-linear summation of induced gamma power across visual subfields. This linearity is particularly apparent in the 40–60 Hz range, where summed gamma curves closely overlay the full-field response in the left/right and top/bottom plots, with minimal deviation. In contrast, the same summation process yields clear subadditivity in the alpha band (8–15 Hz), where the summed alpha suppression (negative deflection in red dashed lines) consistently exceeds that of the full stimulus. This nonlinearity is most evident in the quadrant and octant comparisons, where summed alpha dips to –2.5 to – 4 dB, substantially exceeding the full-field suppression (∼–1 dB), implying nonlinear interaction effects such as saturation or divisive normalization in alpha-band dynamics. The blank condition was subtracted from the summed spectra (blue dotted lines, “Sum(Parts – Blank)”). This adjustment partially restores alignment between summed and full-field alpha responses—particularly in the left/right and top/bottom comparisons—suggesting that shared baseline suppression contributes to alpha subadditivity. However, in more spatially fragmented groupings (e.g., octants), even the blank-corrected sum remains more negative than the full-field alpha response, indicating persistent nonlinear behavior. Thus, gamma summation appears spatially additive and linear, while alpha summation is distinctly nonlinear, only partially mitigated by background correction.

Panel D shows model prediction errors for linear vs. divisive normalization fits across retinotopic groups. For alpha-band power (left), linear fits yield high errors—especially for octants (>2.8) and quadrants (∼1.5)—reflecting the subadditivity seen in Panel C. Divisive normalization sharply reduces error across all groupings, indicating better fit for alpha dynamics. With blank subtraction (center), linear error decreases but remains higher than divisive, particularly for octants and fovea/periphery, suggesting persistent nonlinearity. In contrast, gamma-band fits (right) favor the linear model, which shows minimal error (<0.01) across all comparisons. Divisive normalization overestimates, with error ∼0.06–0.09. These results confirm that gamma power summates linearly, while alpha suppression follows a divisive pattern, only partly corrected by baseline subtraction.

Panel A shows the full set of 16 oriented grating stimuli (Triggers 41–56), spaced at 22.5° intervals from 0° (vertical) to 360°, covering cardinal, oblique, and intermediate directions. These stimuli served as the basis for comparing evoked spectral responses across orientations. Panel B presents results of a one-way repeated measures ANOVA (across stimulus types 41–56), identifying frequency–time windows with significant modulation. The F-statistic map (top left) and corresponding –log10(p) values (top right) show orientation-sensitive differences in the sustained narrowband 40–80 Hz gamma range between ∼0.25–3 s post-stimulus but not alpha. Cluster-based permutation tests (bottom left) revealed a significant gamma-band cluster (p < 0.01), indicating reliable orientation sensitivity in this band. The bar graph (bottom right) summarizes mean gamma-band ERSPs within this cluster to all orientations, showing the highest induced responses for Triggers 51 (112.5°), 50 (135°), and 43 (292.5°), with significantly lower gamma for cardinal orientations (especially horizontal).

Panel C quantifies power modulation in the 0–3 s post-stimulus interval for stimuli grouped according to orientation and drift direction. Gamma power (top left) showed a clear orientation dependence, with strongest responses to oblique orientations (e.g., 112.5°, 135°), and weakest to cardinal (0°, 180°). Alpha power (top right) displayed an inverse suppression profile, strongest at cardinal orientations and in particular horizontal. This relationship was quantified in the scatter plot (bottom left), with a paradoxically near-perfect *positive* correlation between gamma and alpha (r = 0.94, p < 0.005) across orientations (as opposed to the inverse correlation observed for retinotopy). Directional drift grouping (horizontal, vertical, left, right) showed the opposite trend (r = –0.88, p = 0.123), with higher gamma amplitudes and greater alpha suppression for rightward and downward drifting gratings. The lower-right matrix shows frequency–frequency correlations among evoked power spectra based on all stimulus groupings, with a weak positive band around 40–90 Hz (gamma), and weak, non-significant anti-correlation with the 8–12 Hz alpha band, and strong agreement between alpha and beta frequencies.

Panel D explores orientation and directional effects using pairwise t-tests. For orientation (left column), alpha power differences did not reach significance, but trends were observed for oblique vs cardinal angles (e.g., vertical ±22.5° and horizontal ±22.5°) (less alpha suppression for oblique) and in particular for horizontal vs oblique angles (greater alpha suppression for horizontal angles). Unsurprisingly, gamma was more strongly tuned to orientation (bottom left), with significantly higher power for oblique versus cardinal (p < 0.01) and significantly reduced power in horizontal gratings (p < 0.01). Directional comparisons (right column) showed weaker overall effects, with no obvious trends for the alpha range but a significantly increased gamma response to rightward vs leftward gratings (p = 0.013) and a trend towards reduced gamma for up/down drifting gratings vs right-drifting gratins (p = 0.085 and 0.151 respectively). These findings indicate that induced power is more sensitive to absolute orientation than general stimulus direction, with oblique stimuli consistently evoking stronger gamma and but weaker alpha suppression.

## Discussion

Our findings demonstrate a striking dissociation between gamma and alpha rhythmic responses in human visual cortex. Gamma oscillations were highly localized and feature-tuned, summating approximately linearly across distinct stimulus subregions, whereas alpha oscillations were broadly distributed, only weakly selective for stimulus features, and showed subadditive suppressive effects consistent with divisive normalization. This dissociation supports the view that gamma and alpha are generated by distinct neural circuits with complementary roles, rather than being merely inverse reflections of a single process.

### Gamma reflects localized excitation and linear field summation

In our data, gamma-band responses were tightly retinotopic and summed linearly when separate visual patches were stimulated, consistent with primate findings that gamma arises within local cortical networks and reflects the sum of local excitatory drives [12,13]. Non-overlapping stimuli produced additive gamma responses, implying minimal long-range interaction at the generation stage. Gamma also showed robust feature tuning, including orientation and motion direction, with a reliable bias toward non-cardinal orientations—mirroring the “oblique effect” reported in V1 population activity [25–30]. This contradicts invasive studies show higher density of cardinal-preferring neurons [26,27] and perceptual salience of these orientations [28,29], but inverse-oblique effects have been noted recently in EEG gamma [31] as well as visual perception experiments [30].

The linearity we observed matches primate LFP results showing that spatially distant patches generate gamma independently, with saturation only when local inhibitory feedback engages [12,13]. Under our conditions, non-overlapping cortical regions oscillated autonomously at gamma frequencies, indicating gamma power scaled with summed local activity until interactions or saturation occurred. This supports models in which gamma amplitude reflects local excitatory drive within recurrent E–I loops.

We also found evidence for a transition from local to global gamma with large, contiguous stimulation. Foveal and full-field conditions produced disproportionately high gamma power, consistent with macaque studies showing small stimuli evoke local gamma, whereas large stimuli synchronize distant sites into a coherent network oscillation with shared orientation preference [9,13,14]. Murty et al. [14] further showed that beyond a critical size, a second, slower gamma component emerges, greatly boosting total gamma power. This secondary peak at ∼30 Hz. was observed in our data only after subtracting out baseline responses, suggesting that lower frequency suppression can ‘bleed into’ the upper frequencies, masking mid-frequency gamma responses in scalp data. Overall, scalp gamma signals were strongest relative to alpha suppression when large portions of retinotopic cortex oscillated in phase. This suggests EEG/MEG gamma emphasizes the global component of cortical gamma, scaling super-linearly when local generators synchronize. Such network locking explains the reproducibility of gamma frequency and amplitude across cortex and reinforces the view that narrowband gamma indexes coherent cortical excitation, bridging invasive animal studies and human EEG.

### Alpha reflects broad suppression and divisive normalization

Unlike gamma, alpha-band activity showed weak spatial and feature selectivity but was strongly modulated by stimulus presence or expectation. Alpha power dropped sharply whenever a stimulus was anticipated or present, supporting its role as an inverse index of cortical engagement. Enlarging or adding stimuli produced subadditive, saturating suppression: once alpha was already reduced by one patch, a second in a different location caused only a small further drop. This pattern matches divisive normalization models [15,17], where inhibitory feedback scales down additional responses once a certain drive level is reached. Our results suggest alpha indexes this gain-control process, in which overlapping neural populations share an inhibitory signal that limits combined responses. Human ECoG findings similarly show alpha correlating with surround suppression [16], supporting the idea that alpha “tracks” cortical normalization. In effect, gamma summed excitations while alpha divided them, highlighting an antagonistic complementarity.

Mechanistically, alpha’s lack of strong feature tuning and saturating profile suggest origins in global or downstream networks. Physiological evidence points to generation in higher visual areas and thalamocortical circuits linked to feedback [20,21]: in monkey V1, alpha arises mainly in layers 5/6, propagates from higher to lower areas, and disappears when feedback is blocked, while gamma persists. Thalamocortical models show a similar 8–12 Hz “idling” state driven by cortex–thalamus–reticular nucleus loops [20], providing rhythmic inhibition until stronger input arrives. We also observed anticipatory alpha suppression—reductions even for ‘blank’ gray screens following crosshair priming—consistent with top-down modulation. Attention studies show alpha decreases at attended locations and increases in ignored ones [22,23], indicating proactive disinhibition to enhance processing. In our “no-stimulus” cue trials, alpha suppression fit this preparatory role. Overall, alpha’s weak feature tuning, broad reach, divisive summation, and context-dependent suppression align with its proposed role as an inhibitory gain-control signal that dynamically regulates cortical excitation.

### Antagonistic but configuration-dependent relationship between gamma and alpha

We asked whether gamma and alpha oscillations always trade off in a simple push–pull fashion or if their relationship depends on stimulus context. In general, we found the expected inverse coupling: conditions with strong gamma coincided with stronger alpha suppression, consistent with these rhythms jointly maintaining E/I balance [18,19]. However, the relationship was not uniform across all stimulus manipulations.

During orientation tuning, alpha was strongly suppressed for all gratings relative to blank, with no significant differences between preferred and non-preferred orientations (though we did observe a trend toward stronger suppression for horizontal gratings (p = 0.15). In contrast, gamma power varied sharply with orientation, peaking at oblique angles and showing minima for horizontal orientations. Despite the absence of clear alpha tuning, orientation-averaged alpha and gamma responses were *positively correlated* across orientations (r = 0.94, p < 0.005)—a striking reversal of the robust negative gamma–alpha relationship observed for retinotopic manipulations. The stronger alpha suppression for horizontal gratings may reflect anisotropies in early visual cortex, where horizontal orientations—common in natural scenes—drive larger population responses in V1 [28,29], more effectively disengaging alpha-generating inhibitory networks. In contrast, the global gamma LFP shows reduced responses for horizontals, possibly because gamma in EEG reflects spatial summation of locally coherent oscillations; widespread horizontal activation may recruit more diverse orientation domains with less phase alignment, lowering the net global gamma amplitude despite strong local spiking activity [9,13].

Stimulus location and size also shaped the interplay. Foveal and full-field stimuli evoked the strongest gamma responses but relatively weak alpha suppression, whereas peripheral stimulation produced stronger alpha suppression despite weaker gamma. This likely reflects differences in spatial summation: foveal and full field gratings activate a larger, contiguous retinotopic patch, increasing the chance for phase-aligned local generators to sum into strong *global* gamma, while peripheral stimuli drive more spatially dispersed regions with lower cross-site coherence, limiting global gamma even when alpha suppression is pronounced.

### Implications for oscillation theories and cross-species comparisons

The dissociation in gamma and alpha tuning supports models assigning these rhythms to distinct circuit mechanisms: gamma arising from local recurrent E–I networks under feedforward drive (PING), alpha from feedback and modulatory circuits imposing inhibitory control [20,21,24]. This fits hierarchical frameworks like predictive coding, where alpha/beta carry top-down expectations that suppress predictable inputs and gamma carries bottom-up prediction errors. Our finding that alpha can divide down inputs that gamma sums echoes this predictive inhibition. It also aligns with “communication through coherence” [18] and “gating by inhibition” [19] models: gamma synchrony aligns high-excitability phases to open communication channels, while alpha increases inhibition to close them. Large, coherent gamma in low-alpha states (as with large stimuli) likely facilitates inter-areal visual communication; high-alpha states (no-stimulus or surround normalization) decouple networks and reduce throughput. Attention exploits this dual mechanism by lowering alpha and boosting gamma in task-relevant regions [22,23], selectively amplifying some inputs while suppressing others.

Our results link human noninvasive measures to invasive animal data. Human EEG gamma shows feature tuning and spatial summation akin to monkey LFP gamma [9,12– 14], supporting its value for inferring circuit properties, particularly large-scale synchrony. Alpha’s broad suppression and context sensitivity match monkey electrophysiology and human MEG [16,20], reinforcing its role in normalization. Some anisotropies, like the cardinal–oblique bias, are stronger in human population signals than in macaque single units [25–30]; our gamma results capture this population-level bias, suggesting gamma is sensitive to ensemble-scale tuning. Alpha’s lack of significant orientation bias may reflect greater variability in its pre-stimulus baseline, as suppression depends strongly on the starting level of alpha power rather than the specific stimulus features. Together, our findings strengthen a unifying cross-species view: gamma reflects localized, feature-driven excitation and coherent communication; alpha reflects diffuse, feedback-mediated inhibition and competitive normalization. These complementary rhythms jointly regulate information flow through the visual hierarchy.

## Conclusion

Our results reveal a clear dissociation between gamma and alpha rhythms in human visual cortex: gamma was strongly localized, feature-tuned, and summed linearly across non-overlapping regions, whereas alpha was broadly distributed, weakly feature-selective, and showed divisive normalization. Global EEG gamma was maximized when large, contiguous retinotopic regions oscillated coherently (e.g., foveal stimulation), while alpha suppression depended more on pre-stimulus baseline and could be strong even with weak gamma, as in peripheral stimulation. Orientation tuning revealed an unexpected positive alpha–gamma correlation, likely reflecting shared dependence on local excitatory drive, with horizontal gratings producing deeper alpha suppression but weaker global gamma due to reduced cross-site phase alignment. These findings support distinct circuit origins—gamma from local recurrent E–I loops under feedforward drive, alpha from feedback-mediated inhibitory control—and highlight how their interplay depends on stimulus size, location, and feature content. Future experiments could test the mechanisms behind this dissociation by pairing the paradigm with manipulations of attention, expectation, or visual context (e.g., surround suppression) to modulate feedback signals. Applying the same stimuli under varied visual load or task demands, and combining them with source-resolved MEG/EEG or invasive recordings, could more precisely link scalp measures to laminar-and circuit-level generators.

## References

1. Berger, H. (1929). Über das Elektrenkephalogramm des Menschen. Archiv für Psychiatrie und Nervenkrankheiten, 87, 527–570.

2. Adrian, E. D., & Matthews, B. H. C. (1934). The Berger rhythm: Potential changes from the occipital lobes in man. Brain, 57(4), 355–385.

3. Gray, C. M., & Singer, W. (1989). Stimulus-specific neuronal oscillations in orientation columns of cat visual cortex. Proceedings of the National Academy of Sciences, 86(5), 1698–1702.

4. Tallon-Baudry, C., & Bertrand, O. (1999). Oscillatory gamma activity in humans and its role in object representation. Trends in Cognitive Sciences, 3(4), 151–162.

5. Frien, A., Eckhorn, R., Bauer, R., Woelbern, T., & Gabriel, A. (2000). Fast oscillations display sharper orientation tuning than slower components of the same recordings in striate cortex of the awake monkey. European Journal of Neuroscience, 12(4), 1453–1465.

6. Gieselmann, M. A., & Thiele, A. (2008). Comparison of spatial integration and surround suppression characteristics in spiking activity and the local field potential in macaque V1. Journal of Neurophysiology, 99(4), 1857–1872.

7. van Kerkoerle, T., Self, M. W., Dagnino, B., Gariel-Mathis, M. A., Poort, J., van der Togt, C., & Roelfsema, P. R. (2014). Alpha and gamma oscillations characterize feedback and feedforward processing in monkey visual cortex. Proceedings of the National Academy of Sciences, 111(40), 14332–14341.

8. Michalareas, G., Vezoli, J., van Pelt, S., Schoffelen, J. M., Kennedy, H., & Fries, P. (2016). Alpha-beta and gamma rhythms subserve feedback and feedforward influences among human visual cortical areas. Neuron, 89(2), 384–397.

9. Jia, X., Smith, M. A., & Kohn, A. (2011). Stimulus selectivity and spatial coherence of gamma components of the macaque V1 local field potential. Journal of Neuroscience, 31(25), 9351–9360.

10. Murty, D. V. P. S., Shirhatti, V., Ravishankar, P., & Ray, S. (2018). Large visual stimuli induce two distinct gamma oscillations in primate visual cortex. Journal of Neuroscience, 38(11), 2730–2744.

11. Foxe, J. J., & Snyder, A. C. (2011). The role of alpha-band brain oscillations as a sensory suppression mechanism during selective attention. Frontiers in Psychology, 2, 154

12. Gieselmann, M. A., & Thiele, A. (2008). Comparison of spatial integration and surround suppression characteristics in spiking activity and the local field potential in macaque V1. Journal of Neurophysiology, 99(4), 1857–1872.

13. Jia, X., Smith, M. A., & Kohn, A. (2011). Stimulus selectivity and spatial coherence of gamma components of the macaque V1 local field potential. Journal of Neuroscience, 31(25), 9351–9360.

14. Murty, D. V. P. S., et al. (2018). Extensive gamma oscillations and neuronal coherence in macaque V1 cortex in response to naturalistic image sequences. Journal of Neurophysiology, 119(1), 333–345.

15. Carandini, M., & Heeger, D. J. (2012). Normalization as a canonical neural computation. Nature Reviews Neuroscience, 13(1), 51–62.

16. Murray, M. M., et al. (2020). Alpha oscillations reflect spatial suppression in visual cortex. NeuroImage, 207, 116350.

17. Wainwright, M. J., Schwartz, O., & Simoncelli, E. P. (2002). Natural image statistics and divisive normalization: Modeling nonlinearity and adaptation in cortical neurons. Nature Neuroscience, 5(8), 803–821.

18. Fries, P. (2015). Rhythms for cognition: Communication through coherence. Neuron, 88(1), 220–235.

19. Jensen, O., & Mazaheri, A. (2010). Shaping functional architecture by oscillatory alpha activity: Gating by inhibition. European Journal of Neuroscience, 31(12), 2244–2252.

20. Bollimunta, A., et al. (2011). Neuronal mechanisms and attentional modulation of corticothalamic alpha oscillations. Journal of Neuroscience, 31(13), 4935– 4943.

21. Self, M. W., et al. (2017). The role of feedback in shaping the selectivity of cortical neurons. Journal of Neuroscience, 37(22), 5408–5418.

22. Thut, G., Nietzel, A., Brandt, S. A., & Pascual-Leone, A. (2006). Alpha-band electroencephalographic activity over occipital cortex indexes visuospatial attention bias and predicts visual target detection. Journal of Neuroscience, 26(37), 9494–9502.

23. Mathewson, K. E., et al. (2011). Power modulations in alpha and beta oscillations predict behavioral performance in visual spatial attention. Journal of Cognitive Neuroscience, 23(9), 2405–2416.

24. Bastos, A. M., et al. (2015). Visual areas exert feedforward and feedback influences through distinct frequency channels. Neuron, 85(2), 390–401.

25. Furmanski, C. S., & Engel, S. A. (2000). An oblique effect in human primary visual cortex. Nature Neuroscience, 3(6), 535–536.

26. Li, B., Peterson, M. R., & Freeman, R. D. (2003). Oblique effect: a neural basis in the visual cortex. Journal of Neurophysiology, 90(1), 204–217.

27. Wang, Y., et al. (2009). Contrast independence of cardinal preference: Stable oblique effect in orientation maps of ferret visual cortex. European Journal of Neuroscience, 29(6), 1258–1270.

28. Girshick, A. R., Landy, M. S., & Simoncelli, E. P. (2011). Cardinal rules: Visual orientation perception reflects knowledge of environmental statistics. Nature Neuroscience, 14(7), 926–932.

29. Essock, E. A., DeFord, J. K., Hansen, B. C., & Sinai, M. J. (2003). Oblique stimuli are seen best (not worst!) in naturalistic broad-band stimuli: A horizontal effect. Vision Research, 43(12), 1329–1335.

30. Wilson, H. R., Loffler, G., Wilkinson, F., & Thistlethwaite, W. A. (2001). An inverse oblique effect in human vision. Vision Research, 41(13), 1749–1753.

31. Butler, R., Mierzwinski, G.LW., Bernier, P.LM., Descoteaux, M., Gilbert, G., & Whittingstall, K. (2020). Neurophysiological basis of contrast-dependent BOLD orientation tuning. NeuroImage, 206, 116323.

